# Mps1 expression is critical for chromosome orientation and segregation in yeast with high ploidy

**DOI:** 10.1101/2020.03.26.006387

**Authors:** Ashlea Sartin, Madeline Gish, Jillian Harsha, Dawson Haworth, Rebecca LaVictoire, Régis E Meyer

## Abstract

In aneuploid cancer cells, the chromosome segregation apparatus is sensitive to increased chromosome number. The conserved protein kinase, Mps1, is a critical actor of this machinery, orienting the chromosomes properly on the spindle. Abnormally high levels of this kinase have been found in tumors with elevated chromosome number. However, it remains unclear, mechanistically, if and how cells with higher ploidy become dependent upon increased Mps1 levels. To answer these questions, we explored Mps1 dependence in yeast cells with increased sets of chromosomes. We discovered that having more chromosomes affects the ability of cells to orient chromosomes properly. The cells with increased numbers of chromosomes are particularly sensitive to the reduction of Mps1 activity. In *mps1* loss of function mutants, cells display an extended prometaphase with a longer spindle and a delay in orienting properly the chromosomes. Altogether, our results suggest that increased numbers of chromosomes render cells more dependent on Mps1 for orienting chromosomes on the spindle. The phenomenon described here may be relevant in understanding why hyperdiploid cancer cells become excessively reliant on high Mps1 expression for successful chromosome segregation.

**Author summary:** Most cells in solid tumors usually carry far more chromosomes than normal cells. Losing or gaining chromosomes during cell division can lead to aneuploidy (an abnormal number of chromosomes), cancer, and other diseases. Mps1 is a master regulator of cell division that is critical to keep the correct number chromosomes in each daughter cell. This master regulator has been shown to target and affect the function of various actors involved in cell division. Abnormally high levels of this master regulator are found in tumors with elevated chromosome numbers. The high levels of this regulator appear to be protecting these tumor cells. To answer if and how cells with higher ploidy become so dependent of Mps1, we generated yeast cells with increased set of chromosomes. Here, we report that cells with elevated chromosome number are particularly sensitive to the reduction of Mps1 level. In cells with higher ploidy and reduced level of Mps1, the progression during cell division is delayed. In the mutant cells, their ability to properly orient and segregate their chromosomes on the spindle is greatly reduced.

## Introduction

Maintaining error-free chromosome segregation during the cell cycle is critical for normal cell viability. If any aspect of the chromosome segregation process goes wrong, it can lead to aneuploidy which is one of the hallmarks of cancers associated with high risk of tumorigenesis [1]. In order to avoid aneuploidy during mitosis, chromosomes must attach to spindle fibers (microtubules) emanating from opposite poles of the spindle [2–4]. Those connections, also known as bipolar attachment, are mainly supported by a structure called the kinetochore [5]. This proteinacous structure formed by 70-100 different proteins allows the connection between the sister chromatids and the microtubules. Due to the stochastic nature of kinetochore-microtubule associations at early stages of mitosis (prometaphase), many chromosomes initially fail to establish bipolar attachments. Instead, most of the sister kinetochores are initially improperly attached by either being un-attached, having a single kinetochore attached to both poles or having both sister kinetochores attached to the same pole. To correct those erroneous attachments that are detrimental for proper partitioning of the genome, cells rely on a signaling pathway that supports a strict regulation of cell cycle progression. This process known as the spindle assembly checkpoint (SAC) coordinates and facilitates the formation of bipolar attachments [6]. The SAC, which is based on the kinetochore, senses erroneous attachments and delays anaphase onset until all kinetochores are exhibiting bipolar attachment. It’s only when the SAC is fully satisfied that the sister chromatids can segregate into the daughter cells.

In aneuploid cancer cells and in tetraploid cells, whose formation is considered to be one of the major pathways to the formation of aneuploid cells, the chromosome segregation apparatus is sensitive to the increased of chromosome number. This can be observed by an increased level of chromosomal instability (CIN) and an increased dependency on components of the chromosome segregation apparatus [7, 8]. For this reason, targeting those components has become an emerging strategy to preferentially kill cancer cells [9].

The conserved kinase Mps1 is one of the components of the chromosome segregation apparatus [10–12]. This kinase is involved in multiple pathways during the cell cycle: Duplicating the microtubule-organizing center (Spindle pole body in yeast, centrosome in mammals), controlling the SAC, and re-orienting chromosomes on the spindle to promote the formation of bipolar attachments. Interestingly, high levels of this kinase correlate with the high degrees of aneuploidy in cancer cells and tumor aggressiveness. Further, increased levels of Mps1 contributes to the survival of aneuploid cancer cells [13]. Similarly, higher levels of Mps1 are also critical for cells with elevated ploidy such as tetraploid cells [14]. For these reasons, the development anti-Mps1 compounds [15, 16] has become a potential cancer therapy (reviewed in [17]). However, despite this therapeutic potential, it still remains unclear, mechanistically, how cells with elevated chromosome number such as aneuploid or tetraploid cells become so dependent upon Mps1 levels. One hypothesis is that having extra-chromosomes places a burden on the chromosome segregation machinery. By this model, excess Mps1 becomes necessary to properly segregate the higher number of chromosomes into daughter tumor cells. Previously, it has been shown that cells with increased ploidy are defective in their ability to promote bipolar attachment due to an inadequate upscaling of spindle geometry/size [18]. But Mps1 has not been evaluated in this model.

To determine if and how cells with increased chromosome numbers become dependent on Mps1, we explored Mps1 dependence in yeast cells with a range of ploidy. By using incomplete inactivation of Mps1, we demonstrated that *mps1* mutants exhibit ploidy-specific lethality. We found that *mps1* mutant cells display extended metaphases with and produce longer spindles. In wild-type cells, we found that the initial attachment of kinetochores to the nascent spindle is not profoundly different in haploid and tetraploid cells. Those initial erroneous attachments can be corrected into proper attachments but the efficiency depends on the spindle length. The longer the spindle, the less efficient is the conversion of incorrect to correct attachments. As ploidy increases, this conversion is more affected. Interestingly, in *mps1* mutant cells, the ability to promote the formation of correct attachments before the anaphase onset is greatly compromised.

## Results

### Ploidy-specific sensitivity and lethality are associated with Mps1 reduced level

To identify which cellular event(s) in cells with increased ploidy are the most compromised when Mps1 activity is reduced, we created a range of inactivation for this kinase. This was done using combinations of the wild-type allele (*MPS1*) and two different mutated versions (*mps1Δ* and *mps1-R170S*). We either used the deletion to fully remove *MPS1* (*mps1Δ*) or a partial loss of function (*mps1-R170S*). The *mps1-R170S* mutation is located in a domain previously defined as required for the spindle checkpoint and re-orienting the chromosome but without defects in the SPB duplication [19]. Cells bearing the *mps1-R170S* allele are perfectly viable and without noticeable defects in SPB duplication [20]. However, the mutant exhibits defects in the spindle checkpoint and in meiotic chromosome segregation [20]. To prevent the accumulation of suppressor mutations, in strains bearing the mutated versions of *MPS1*, we introduced into the cells a shuffling plasmid containing the *TRP1* yeast marker and a copy of *MPS1-as1*. The *MPS1-as1* allele produces an analog-sensitive version of Mps1 that behaves like a wild-type copy unless inhibitor (such as 1-NMPP1) are added to the medium. The shuffling plasmid was either selected by transferring cells on plate without tryptophan (SC-TRP) or counter-selected on plate with FAA which is toxic for *Trp*+ Cells (FAA). This allow us to assess the requirement of the strains for the *MPS1-as1* copy.

For all the ploidies tested, cells carrying one copy of *MPS1* per genome were viable (S1 Fig). Conversely all *mps1Δ* mutant cells were unable to grow (S1 Fig). The *mps1-R170S* mutant cells were viable for all the ploidies tested (S1 Fig). To optimize the extent of the reduction of Mps1 activity in diploid and tetraploid cells, we used combinations of three different alleles: *MPS1*, *mps1Δ* and *mps1-R170S* (Fig 1). To test whether yeast, like mammalian cells, are sensitive to reductions in Mps1 activity when they are polyploid, we removed half of the *MPS1* copies in diploid and tetraploid cells by using the deletion allele (*mps1Δ)*. For the remaining copies, we either used one or two copies of the wild-type allele (Respectively *MPS1-1x* in diploid, Fig 1A or *MPS1-2x* in tetraploid, Fig 1B) or the *mps1-R170S* allele (Respectively *mps1-R170S-1x* in diploid, Fig 1A or *mps1-R170S-2x* in tetraploid, Fig 1C). In the diploid, no major defects were observed in *MPS1-1x* mutant (Fig 1A) whereas the *MPS1-2x* tetraploid mutant exhibited reduced viability (Fig 1B). In *mps1-R170S-1X* diploid mutant, we observed a decreased viability (Fig 1A). Interestingly, in *mps1-R170S-2x* tetraploid mutants, the effect was even more drastic as we didn’t observe any viable cells (Fig 1C). Considering this compelling ploidy-specific sensitivity to the reduction of Mps1 activity, we decided to investigate further the origin of this phenotype. We conclude that yeast cells, like mammalian cells, show a ploidy-dependent sensitivity to Mps1 activity.

**Fig 1.**
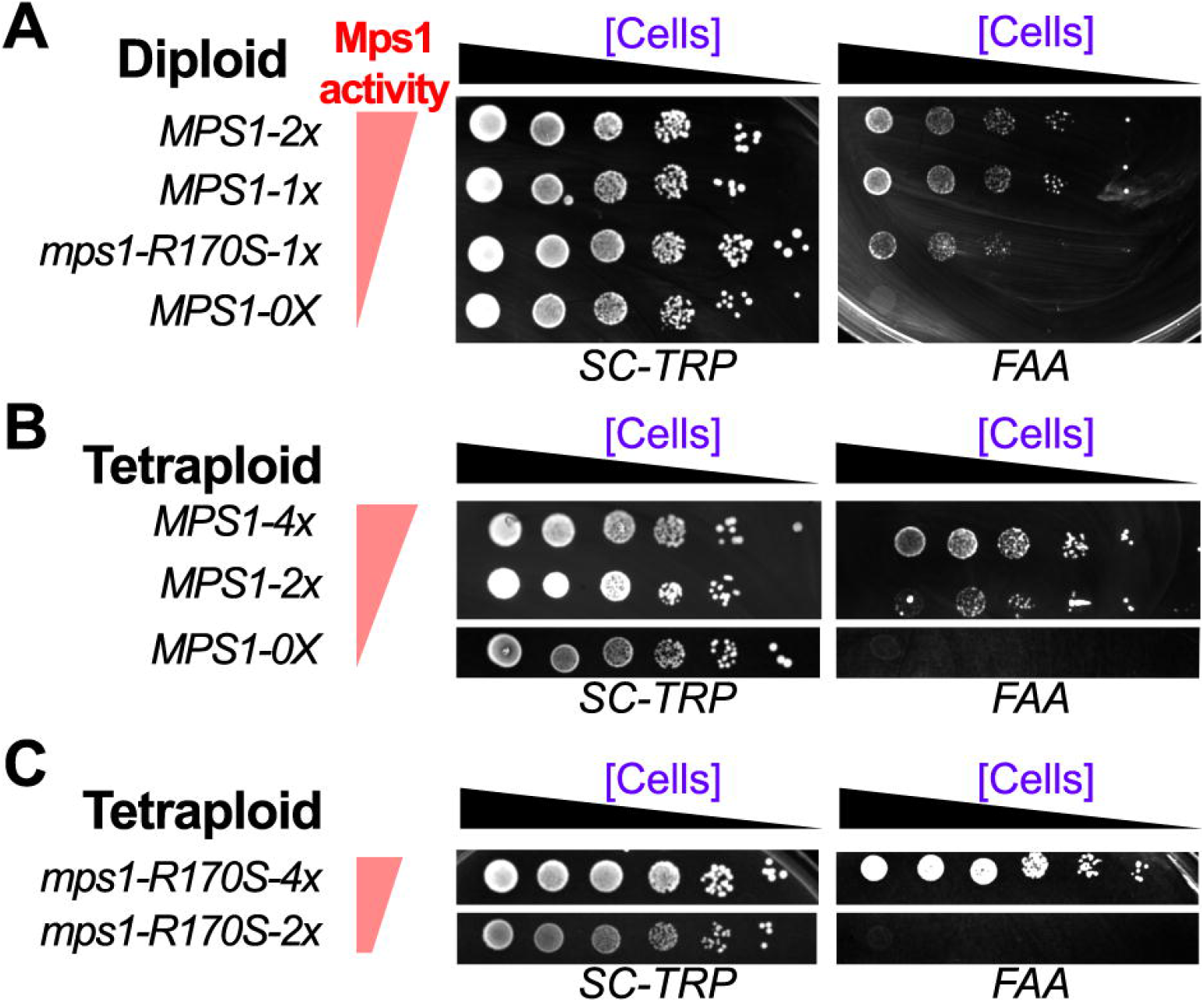
The mps1 mutants display a ploidy-specific sensitivity. (A-C) Ten-fold serial-dilution growth assays. All the indicated strains are carrying a shuffling plasmid expressing *MPS1-as1* and the yeast marker *TRP1*. Cells were originally grown in CM-TRP media and later spotted on the indicated plates. The CM-TRP plates were used to select yeast carrying the shuffling plasmid with the extra copy of Mps1 whereas the FAA plates were used to counter-select it. (A) Diploid cells with either two copies of *MPS1* allele (*MPS1-2x = MPS1/MPS1)*, one copy of *MPS1* allele (*MPS1-1x = MPS1/mps1Δ)*, one copy of *mps1-R170S* allele (*mps1-R170S-1x = mps1-R170S/mps1Δ)* or none of them (*MPS1-0x = mps1Δ/mps1Δ*) were used in this assay (B) Tetraploid cells with either four copies of *MPS1* allele (*MPS1-4x = MPS1/MPS1/MPS1/MPS1*), two copies (*MPS1-2x = MPS1/MPS1/mps1Δ/mps1Δ)* or none (*MPS1-0x = mps1Δ/mps1Δ/mps1Δ/mps1Δ*) were used in this assay. (C) Tetraploid cells with either four copies of *mps1-R170S* allele (*mps1-R170S-4x = mps1-R170S/mps1-R170S/mps1-R170S/mps1-R170S*) or two copies (*mps1-R170S-2x = mps1-R170S/mps1-R170S/mps1Δ/mps1Δ*) were used in this assay.

### Examining the requirements for Mps1 in polyploid cells using live cell imaging

Mps1 is essential for duplicating SPBs, the spindle checkpoint, and promoting the proper attachments of chromosomes to the spindle. To begin to distinguish among these functions as those that are especially critical to disruption in polyploid cells we monitored cell growth using live cell imaging. SPBs were labeled with GFP (Spc29-GFP) allowing the monitoring of SPB duplication and timing of metaphase duration. Cells were placed in microfluidic chambers and imaged for 8 hours. To examine how the reduction in Mps1 activity affects these processes we generated of Mps1 with an auxin degron (*MPS1-AID\**) [21, 22]. This system allows the rapid degradation of the targeted protein in response to auxin addition to the medium. We initially confirmed that tetraploid cells tagged with *MPS1-AID\** were non-viable in the presence of Auxin (S2 Fig). To study the impact of partial Mps1 reduction, we maintained a single wild-type copy of Mps1 (*MPS1-1x*) in all the ploidies, with the remaining copies of the gene either a deletion or the *MPS1-AID\** version. Therefore, by adding auxin to the media, we either removed one copy of the two copies in diploid (*MPS1/mps1-aid\**), two of the three copies in triploid (*MPS1/mps1-aid*/mps1Δ*) or three of the four copies in tetraploid (*MPS1/mps1-aid*/mps1Δ/mps1Δ*).

We initially evaluated the ability of cells with different ploidies to divide and grow inside the microfluidic chambers (S3A Fig). For this purpose, we tracked if a cell can form a bud and/or a spindle after each cell division (S3B Fig). Focusing on the initial cell divisions, we didn’t notice any significant difference in cell viability between ploidies in wild-type cells (4.9% vs. 7.0%, n>43, Fig 2A). However, we noticed that *MPS1-1x* mutant tetraploid cells display a significant increase of dead cells compare to the wild-type cells (40.5%, n=115). This was easily noticeable by the frequent appearance of cells unable to grow (S3B Fig).

**Fig 2.**
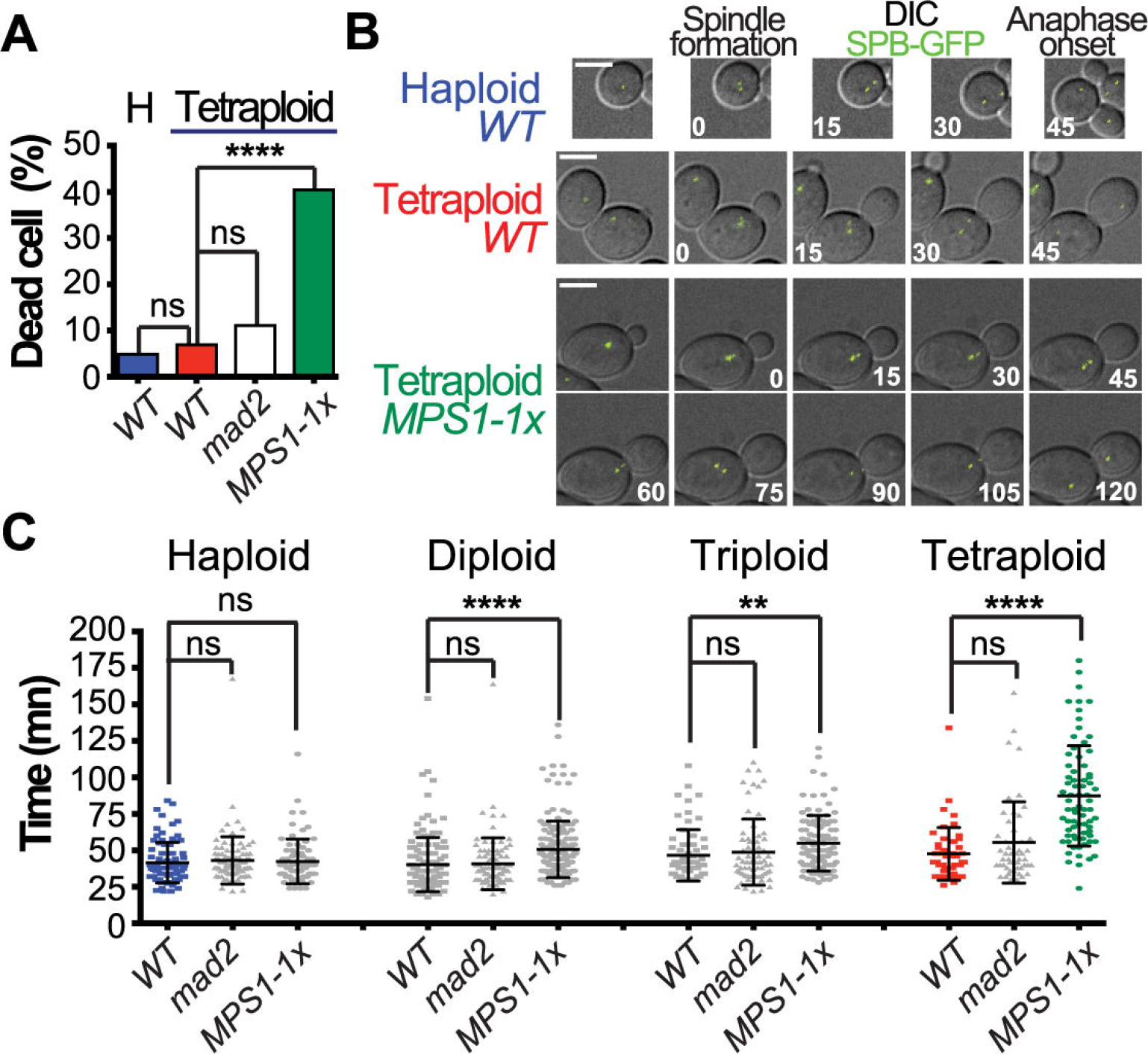
Anaphase onset is delay in mps1 mutants. (A-C) All cells (haploid, diploid, triploids and tetraploid) of the indicated genotype are carrying a single copy of Spc29-GFP. The cells were grown in SD media and observed by a time-lapse movie at 120-150 second intervals for 4-8 hours. The cells are all expressing *P_CUP1-_AFB2*, a F-protein necessary for the degradation in yeast cells of the AID* tagged protein. For the *MPS1-1x* genotype, a single *MPS1* allele remain in all ploidy. The haploid *MPS1-1x* cells retains a single *MPS1* allele. For the diploid *(MPS1/mps1-AID*)*, triploid *(MPS1/mps1-AID*/mps1Δ)* and tetraploid *(MPS1/mps1-AID*/mps1Δ/mps1Δ),* auxin and copper were added to the media to remove the AID* tagged version of Mps1. (A) The proportion of dead cells was determined during the 1^st^ two cell cycles following each cell division (n≥43, see S3 Fig for detail). Asterisk indicated a statistical difference between the wild-type cells and the mutants tested. **** = p < 0.0001, ns=non-significant (Fischer’s exact t test). (B) Representative pictures from wild-type cells (haploid and tetraploid) and *MPS1-1x* mutant tetraploid cells are shown. Bars 5 μm. Indicated time is in minutes. (C) Each indicated genotype was tested in at least three independent experiments. Time from SPB separation to anaphase onset was measured for each cell of the indicated genotype and ploidy. Dot represents one data point collected from time-lapse microscopy. Asterisk indicated a statistical difference between the wild-type cells and the mutants tested ** = p < 0.01, **** = p < 0.0001, ns=non-significant (Unpaired t test). Error bar are the SD.

### Are polyploid cells sensitive to Mps1’s function in spindle pole body duplication?

Investigating the impact of partially removing Mps1 on the ability of cells to divide, we were able to track critical steps of cell division (spindle formation, anaphase) over several consecutive cell cycles. Therefore, this result suggests that the partial removal of Mps1 only affect modestly the SPB duplication process. To define a potential defect, we evaluated the ability of each tetraploid cells to duplicate their SPBs. For this purpose, we used the formation of a bud as a marker of cell cycle progression and tracked if those cells were able to display two distinct SPBs following this event. Confirming our previous results, we observed that the majority of the *MPS1-1x* mutant cells can indeed duplicate their SPBs. However, in comparison to the wild-type tetraploid cells, we noticed a significant proportion of cells unable to do so (S3C Fig). We can’t rule out that the *MPS1-1x* mutant cells just happened to stop growing after forming a bud independently of a defect in duplicating SPB. However, this later result confirms that partially removing Mps1 activity have no or only a modest impact on SPB duplication.

### Are polyploid cells sensitive to Mps1’s function in the spindle checkpoint?

Mps1 acts near the top of the spindle checkpoint by phosphorylating Mad1 and triggering metaphase delay when chromosomes are improperly attached. Prior studies have suggested that polyploid cells are viable when the spindle checkpoint is inactive (*mad2*) suggesting that this is likely not the function of Mps1 that polyploid cells are especially dependent upon. If polyploid cells with reduced Mps1 activity have lost the spindle checkpoint function the prediction is they would have a shortened metaphase. To test this, we measure the duration of metaphase in cells of increasing ploidy, with full or reduced levels of Mps1. As a control we also measured metaphase duration in cells lacking the essential spindle checkpoint gene, *MAD2*. In wild-type cells, the there is a small but significant increase in metaphase duration with increasing ploidy (41.5mn ± 1.5mn in haploid vs. 46.3mn ± 2.2mn in triploid and 47.6mn ± 2.6mn in tetraploid, Fig 2B-C). This increase is consistent with the hypothesis that polyploid cells are confronted with bi-orienting larger numbers of chromosomes and this takes more time. This increase in metaphase duration might be expected to be triggered by a spindle checkpoint delay. However, deleting *MAD2* did not reduce metaphase duration, suggesting either that the longer metaphase in polyploid cells is not due to the checkpoint, or alternatively that *MAD2* has other functions that when removed, slow down metaphase. As expected by previous results in tetraploid cells, we also failed to notice any significant impact of removing spindle checkpoint on cell viability (Fig 2A).

In contrast, reducing Mps1 levels significantly increased the duration of metaphase in a ploidy dependent fashion. In diploid cells, removing half of the Mps1 lead to an increase of 10 minutes in the timing from spindle formation to anaphase comparison when compared to wild-type cells (50.7mn ± 1.9mn vs. 40.3mn ± 1.6mn, Fig 2C). We observed a similar increase following the removal of 2/3 of Mps1 copies in the triploid cells (54.9mn ± 1.9mn vs. 46.6mn ± 2.2mn). Finally, we noticed the most compelling increase when we removed ¾ of the Mps1 copies in tetraploid cells ended-up by doubling the duration of mitosis (87.3mn ± 4.0mn vs. 47.6mn ± 2.5mn, Fig 2C).

### Extended metaphase displays longer spindle

As yeast cells bi-orient their chromosomes in pro-metaphase, the spindle gradually gets longer [23–26]. The spindle is restrained from extending because bi-oriented sister chromatids are held together by cohesins. The gradual spindle lengthening in pro-metaphase could be due to gradual loss of centromeric sister chromatid cohesion [27, 28]. It was shown previously that spindle length is not affected by the changes of ploidy [18]. But this conclusion was drawn by measuring the length of fixed metaphase spindles. By tracking the distance in 3D between the two SPBs, we were able to follow for individual cells their spindle length depending of the amount of time spent in pro-metaphase (See trace for haploid and tetraploid cells in Fig 3A). Comparing haploid and tetraploid cells, we noticed that it is the amount of time spent in pro-metaphase, not the ploidy, that determines spindle length (Fig 3B).

**Fig 3.**
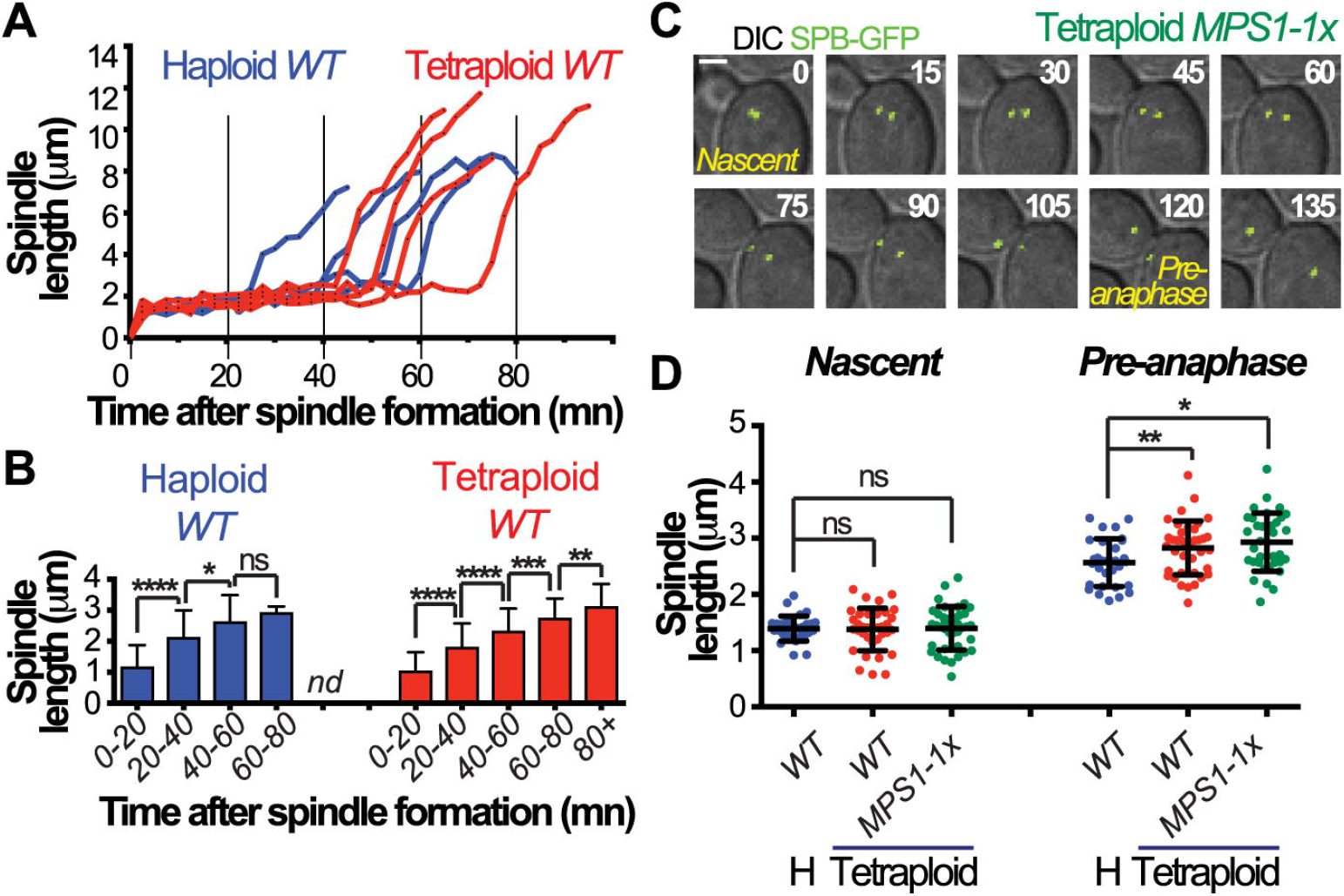
Pre-anaphase spindle get longer when anaphase onset is delay. (A-D) All cells (haploid, diploid, triploids and tetraploid) are carrying a single copy of Spc29-GFP. The cells were grown in SD media and observed by a time-lapse movie at 120-150 second intervals for 4-8 hours. Strain genotypes are as in Fig 2. The spindle length was estimated by measuring in 3D distance between the two SPBs. (A) For each wild-type cell (Haploid and Tetraploid, n≥10), the measurement was done every 150 seconds from prometaphase to the anaphase onset. (B) The graph represents the average of those measurement for the indicated period of time spent in prometaphase. * = p < 0.05, ** = p < 0.01, *** = p < 0.001, **** = p < 0.0001, ns=non-significant (Unpaired t test). Error bar are the SD. (C) Representative pictures from the spindle formation (Nascent spindle) to the anaphase onset in a *MPS1-1x* mutant tetraploid cell are shown. Bars 2 μm. Indicated time is in minutes. (D) For the indicated genotype, the measurement was either done on nascent spindle or just before the spindle elongate (Pre-anaphase spindle). At least 30 cells were counted per genotype. * = p < 0.05, ** = p < 0.01, ns=non-significant (Unpaired t test). Error bar are the SD.

The extended metaphase duration of polyploid cells (Fig 2C) suggests they may actually have longer spindles than haploid cells by the end of metaphase when the last chromosomes are bi-orienting. To test this, we monitored spindle length through pro-metaphase as a function of ploidy and *MPS1* dosage. We demonstrated that when the spindle form, their lengths are not significantly different across ploidies (haploid *WT*: 1.40μm ± 0.04μm, tetraploid *WT*: 1.38μm ± 0.06μm; tetraploid *MPS1-1x*: 1.40μm ± 0.06μm mean ± SEM). However, for spindles just before anaphase onset (pre-anaphase, Fig 2C), cells with higher ploidies have significantly longer spindles (haploid *WT*: 2.56μm ± 0.08μm, tetraploid *WT*: 2.93μm ± 0.09μm, tetraploid *MPS1-1x*: 2.82μm ± 0.08μm, Fig 2D).

### The ability to convert monopolar into bipolar attachments is impacted by both spindle length and the ploidy

Previously, it was determined that polyploid cells with metaphase spindles exhibit a greater proportion of mono-oriented chromosomes than euploid cells with metaphase spindles [18]. However, since these experiments were done with fixed cells, it was unclear if the defect was originating from a defect in the initial attachment and/or in the conversion of the monopolar into bipolar attachment. To address this question, we used live-cell imaging to track chromosome bi-orientation as a function of both spindle length and ploidy. To reveal the movements of chromosomes at higher resolution than in previous experiments, we imaged chromosome behavior at much faster acquisition rates (5 to 10 second intervals) over the course of five minutes capturing a window of chromosome behavior in prometaphase. To reduce acquisition times, the SPBs and the centromere of chromosome I were both tagged with GFP. Chromosome behavior was quantified in cells that had bipolar spindles at the beginning of the imaging window (t=0mn, see Fig 4A). We assigned cells into one of three categories based upon the chromosome behavior. These included the monopolar attachment (sister centromeres together and on one side of the spindle), bipolar attachment with un-separated sister centromeres (revealed by a central position on the spindle) or bi-polar attachment with separated sister centromeres. In haploid, the cells with short spindles (less than 0.75μm) exhibit mostly monopolar attachment and as the spindle gets longer, those attachment are converted into bipolar attachments (Fig 4A). This work is consistent with previous studies showing that sister chromatids usually begin mitosis mono-oriented and become bi-oriented over time while the spindle length gradually increases in pro-metaphase [23–26].

**Fig 4.**
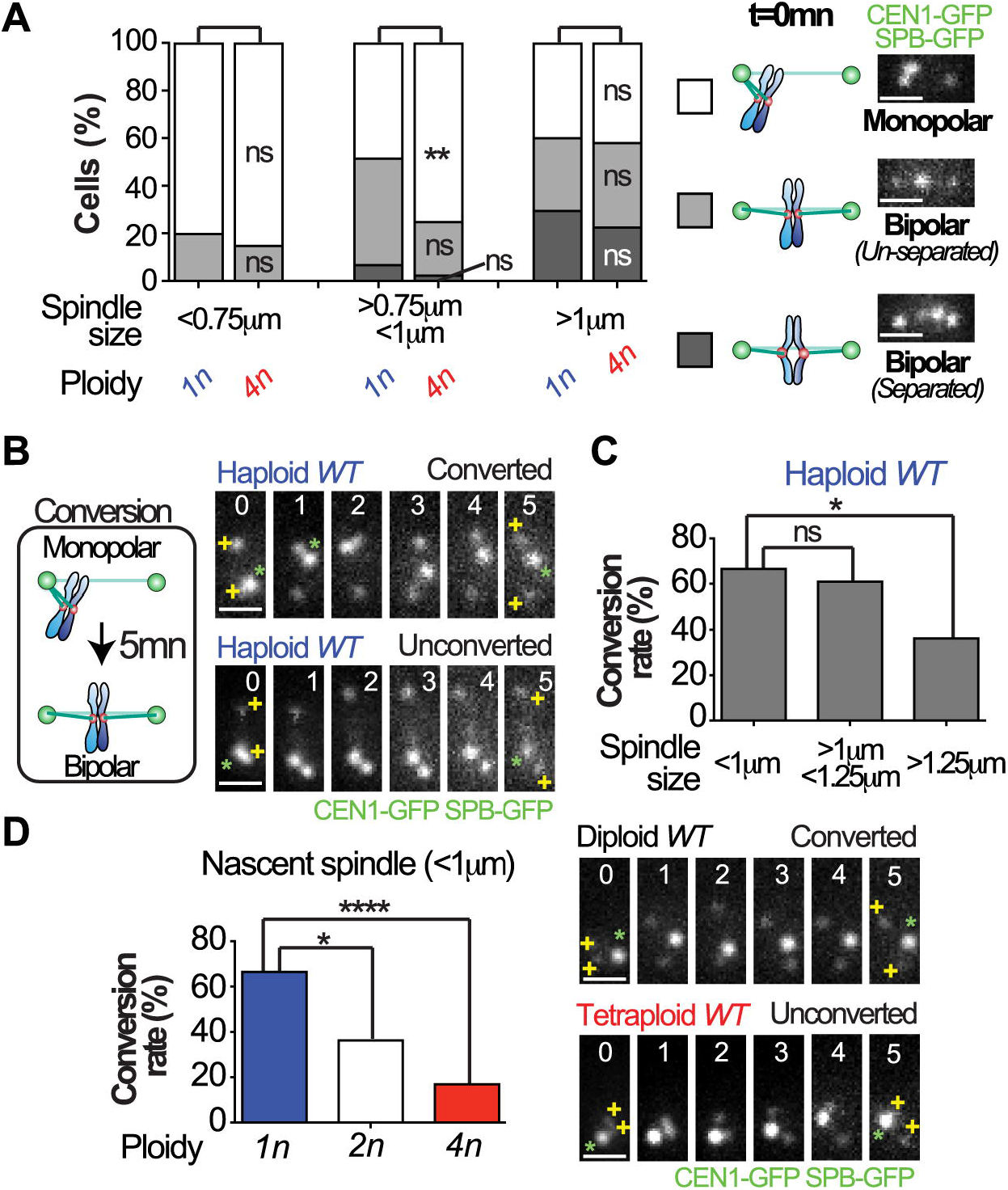
Conversion from monopolar to bipolar attachment is reduced on longer spindle and in higher ploidy. (A-D) All wild-type cells (haploid, diploid and tetraploid) are carrying a single copy of Spc29-GFP and one *CEN1-GFP* tagged chromosome. Cells were observed by time-lapse imaging at 5-10 second intervals for 5 minutes. (A) Cells were placed in categories according to the behavior of *CEN1* at the beginning of the acquisition time (t=0mn). Cells with *CEN1* staying near-by the poles were classified as monopolar attachment (aka syntelic/mono-oriented). Cells with *CEN1* staying in the middle of the spindle were classified as bipolar attachment (aka amphitelic/bi-oriented) with either un-separated or separated sister chromatid (if their splitting was observed). The proportion of each categories depending of the indicated spindle length was estimated for the indicated ploidy. At least 137 cells were counted per ploidy. ** = p < 0.01, ns=non-significant (Fischer’s exact t test). Representative pictures of the type of attachment are shown. Bars 1 μm. (B) The ability of individual cell to convert monopolar to bipolar attachment in 5 minutes was evaluated. Representative pictures of cells able or unable to convert monopolar attachment are shown. The green asterisks represent CEN1 and the yellow plus signs represent the SPBs. Bars 1 μm. Indicated time is in minutes. (C-D) The proportion of cells able to convert in 5mn interval depending of the spindle size in haploid wild-type cells (C) or the indicated ploidy on nascent spindle (D) are shown. At least 20 cells per category of size or ploidy were counted. * = p < 0.05, **** = p < 0.0001, ns=non-significant (Fischer’s exact t test). Representative pictures of cells with nascent spindle able or unable to convert monopolar to bipolar attachment are shown. The green asterisks represent CEN1 and the yellow plus signs represent the SPBs. Bars 1 μm. Indicated time is in minutes.

Tetraploid cells with short (nascent) spindles exhibited similar behavior to haploid cells with short spindles: most of the sister chromatid pairs entered pro-metaphase mono-oriented on the spindle (Fig 4A). However, as cells are progressed through metaphase, the proportion of cells with monopolar spindles decreased more slowly than was seen in haploid cells. This result suggests that the ability of cells to convert monopolar into bipolar attachments is diminished by the number of chromosome present. Therefore, to confirm this idea, we analyzed the ability of a cell to convert a monopolar attachment into a bipolar attachment (separated or un-separated sister chromatid) in 5mn interval (Fig 4B). Among haploid cells, those with short spindles were the most efficient in the conversion and while those with longer spindles reduced the rate of conversion (Fig 4C). Interestingly we noticed that as the ploidy is increasing, the ability to convert monopolar attachment is gradually reduced on the short spindle (Fig 4D). Surprisingly, tetraploid cells exhibit a similarly low rate of conversion independent of the spindle length (S4 Fig).

### Converting monopolar into bipolar attachments is delayed with Mps1 reduced level

Using the same analysis, we investigated whether Mps1 reduction affects the initial attachments, the ability of cells to convert monopolar into bipolar attachments and whether *mps1* mutant cells exhibit ploidy sensitivity in the bi-orientation process. *MPS1-1x* tetraploid mutant cells exhibited an even higher proportion of monopolar attachments on the nascent spindles (Fig 5A). On the longer spindle, the proportion of *MPS1-1x* tetraploid mutant cells exhibiting monopolar attachment is reduced but still much higher than the wild-type tetraploid cells. This result suggests that the reduction of Mps1 activity doesn’t fully abolish the ability of cells to convert mono-polar into bi-polar attachments but instead delay this process. This was confirmed by the lower rate of conversion from monopolar to bipolar attachments in *MPS1-1x* mutant cells compare to the wild-type cells (Fig 5B). To study the impact of this lower rate of conversion on the final attachment of sister chromatid, we investigate how they attach to the spindle just before the anaphase onset (t=−60s, pre-anaphase spindle, Fig 5C). On those spindles, we observed that the haploid cells exhibit a typical bipolar attachment with a high stretching of the sister chromatids (distance>0.5μm). However, we noticed a significant reduction of this type of attachment in tetraploid cells. Instead, the wild-type tetraploid displays an increased proportion of bipolar attachment with a reduced stretching between sister chromatid. The *MPS1-1x* mutant cells appears to be even more affected, exhibiting a high proportion of cells with a monopolar attachment at this stage (Fig 5D). Even do, we only noticed a modest proportion of cell resulting in non-disjunction event in *MPS1-1x* mutant cells (1 of 15 cells), keeping monopolar attachment at this stage can be detrimental for the cells as the re-orientation process is greatly reduced on those longer spindle (see Fig 4C and S4 Fig).

**Fig 5.**
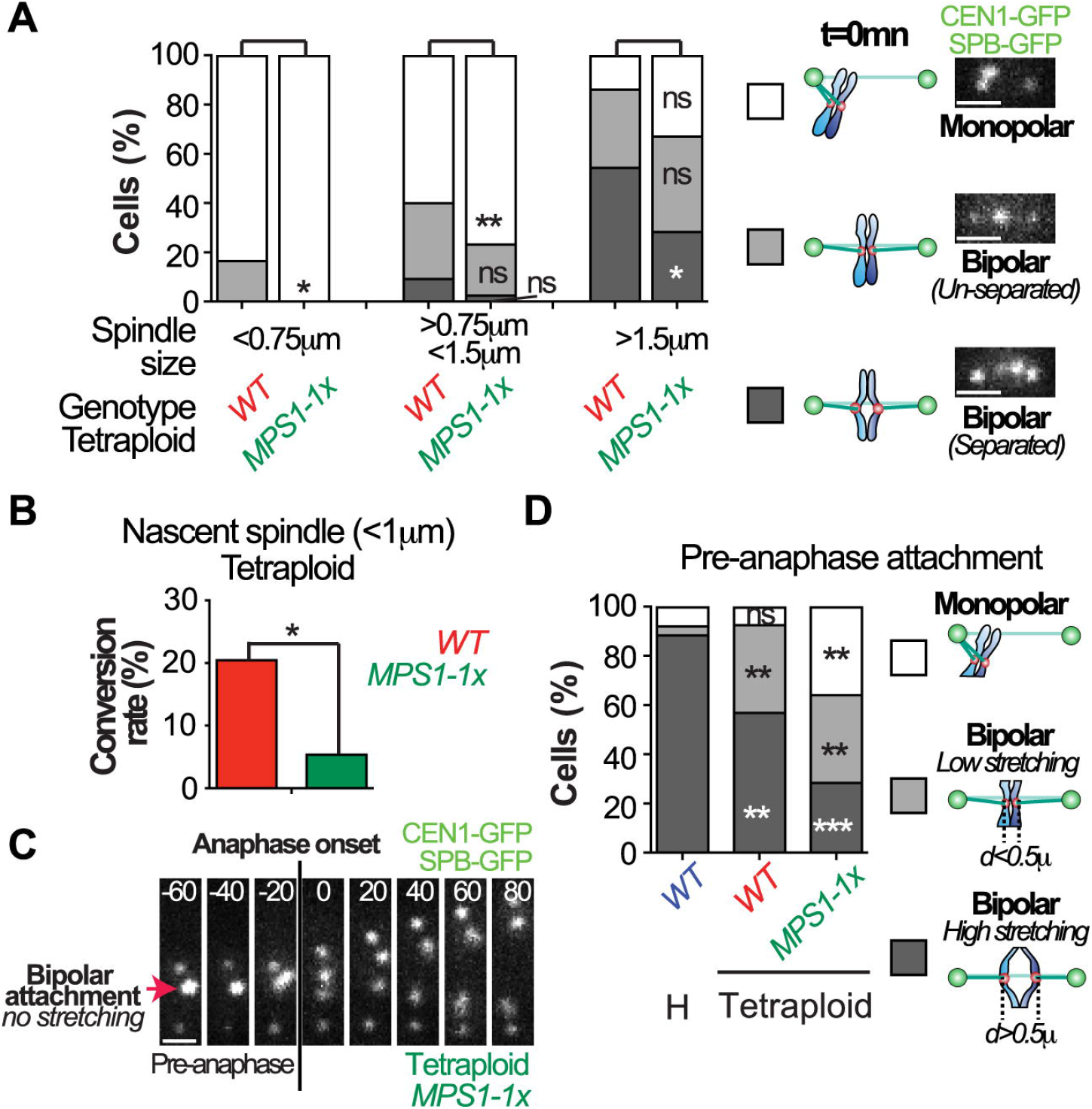
The formation of bipolar attachment is delayed in mps1 mutants. (A-D) All wild-type and *MPS1-1x* mutant cells (haploid and tetraploid) are carrying a single copy of Spc29-GFP and one *CEN1-GFP* tagged chromosome. Cells were observed by time-lapse imaging at 5-10 second intervals for 5 minutes. (A-B) The assays are the same as described in Fig 4. (A) The proportion of each categories depending of the indicated spindle length for the indicated genotype of tetraploid cells is shown. At least 170 cells were counted per genotype. * = p < 0.05, ** = p < 0.01, ns=non-significant (Fischer’s exact t test). (B) The conversion rate of tetraploid cells of the indicated ploidy on nascent spindle (<1μm) was estimated. At least 39 cells per genotype were counted. (C-D) The type of attachment in haploid or tetraploid cells of the indicated genotype was evaluated 1 minute before the anaphase onset. Cells were placed in categories according to *CEN1* behavior at this time (t=−60s). Cells with *CEN1* staying near-by the poles were classified as monopolar attachment. Cells with *CEN1* staying in the middle of the spindle were classified as bipolar attachment with either low stretching (if the distance between the two sister chromatids was lower than 0.5μm) or high stretching (if the distance between the two sister chromatids was higher than 0.5μm). (C) Representative pictures of *MPS1-1x* mutant tetraploid cells displaying a bipolar attachment with low stretching one minute before the anaphase onset are shown. Indicated time is in seconds. Bars 1 μm. (D) The proportion of each categories depending of the indicated ploidy and genotype is shown. At least 14 cells were counted per genotype and ploidy. ** = p < 0.01, *** = p < 0.001, ns=non-significant (Fischer’s exact t test).

## Discussion

The proper segregation of sister chromatids during mitosis is affected by the number of chromosomes. This can be easily seen when the ploidy is increasing such as in budding yeast and mammal tetraploid cells, as they exhibit higher levels of chromosomal instability than euploid cells. One of the weaknesses of polyploid cells is the ability of their chromosome segregation apparatus to adapt to the increase of chromosome number. In support of this idea, it was shown that yeast cells with increased ploidy are particularly sensitive to the inactivation of key actors of the chromosome segregation apparatus [18, 29]. By investigating the ability of chromosome to properly attach to the spindle, it was shown that cells with increased ploidy display more erroneous (= monopolar) attachments on the metaphase spindle than haploid cells. However, it was unclear which of the step(s) that promote the formation of bipolar attachments was affected.

To answer this question, we tracked chromosome re-orientation using live-cell imaging as a function of both ploidy and Mps1 levels. Surprisingly, we discovered that the initial proportion of erroneous attachments on the shortest spindle is close between all the ploidies. The impact of increasing the chromosome number only happens as cells progress into mitosis. Using haploid cells, we demonstrated that the ability of cells to convert erroneous attachment into proper attachments strictly depends on their progression into pro-metaphase. On short spindles (early pro-metaphase), monopolar attachments are rapidly converted into bipolar attachment whereas on long spindles (late pro-metaphase), this ability is greatly reduced. Comparing this process among all the ploidies, revealed that the more chromosomes are present into the cells the more they the cells struggle to re-orient them. Therefore, the main difference between ploidies is not on the proportion of initial erroneous attachment but more on how those attachment can be converted into proper bipolar attachment.

To find why cells are so dependent of Mps1 when the chromosome number increases, we reduced Mps1 levels and studied its impact on chromosome segregation. We identified four striking features in *mps1* mutant cells: 1) The transition from metaphase to anaphase is delayed 2) The spindle length is extended during the stage just before the anaphase onset (pre-anaphase) 3) The conversion from monopolar to bipolar attachment is delayed 4) Cells are showing an increased sensitivity to inactivation at higher ploidy. Hauf and colleagues described a compelling model that can account for the defect that we observed in *mps1* mutant cells and higher ploidy. They noticed an impact on re-orienting chromosome when spindle length was increased [30]. They proposed that two processes are acting in parallel to get mono-oriented kinetochores from the pole to the mid-zone and termed these the direct and indirect routes. In the direct route, microtubules from one pole reach across the spindle to connect to mono-oriented kinetochores, and drag them back towards the mid-zone (Fig 6). In the indirect route, kinetochores can slide along microtubules that emanate from the pole where they are clustered, toward the plus ends. It was later found this kind of movement, mediated by motor-plus end proteins can be observed in mammalian cells and fission yeast [31, 32]. It’s is likely happening in budding yeast, but has not been reported.

Together, our data support the following model that can explain why the chromosome segregation apparatus is sensitive to the increase of chromosome number and why cells become so depend on Mps1 action. On short spindles, monopolar attachments can be easily converted into bipolar attachments using the direct route (Fig 6). Since the distance for the microtubules emanating from one pole to reach across the spindle to the mono-oriented kinetochores is short, it’s easy to promote the formation of bi-orientation of sister chromatid. However, on longer spindles, we propose this process is compromised, and needs an additional mechanism such the indirect route that can slide chromosome on the microtubule to reach microtubule plus-ends from the opposite pole. By this model, as the number of chromosomes increases, the time to bi-orient the chromosomes increases which leads to longer spindles. The longer spindles are less efficient at bi-orienting the remaining mono-oriented chromosomes, and processes that require Mps1 to assist in bi-orientation. Thus, rendering polyploid cells especially vulnerable to reductions in this kinase. Learning mechanistic details of how Mps1 helps chromosomes become bi-oriented will assist in revealing in more detail the vulnerability of highly aneuploid tumor cells to reductions in Mps1 activity.

## Materials and Methods

### Yeast strains and culture conditions

All strains are derivatives of two strains termed X and Y described previously [33]. We used standard yeast culture methods [34]. The yeast strains used in this study are lister in Table S1-3. For the long-term time-lapse microscopy, the diploid, triploid and tetraploid *MPS1-1x* mutants, auxin (0.5mM, Sigma Aldrich I5148-10G) and copper (200μM, Sigma Aldrich 451657-10G) were added after cells were loaded into the chamber. For the short-term time-lapse microscopy with tetraploid *MPS1-1x* mutants, auxin (0.5mM, Sigma Aldrich I5148-10G) and copper (200μM) were added 30mn before cells were concentrated on coverslips. For the serial-dilution assay, auxin (0.5mM) was added to YPAD plate.

### Generation of triploid and tetraploid cells

The tetraploid cells were generated by mating diploid strains of opposite mating types (Diploid *Mata*/*Mata* with diploid *Matα*/*Matα*). The triploid cells were generated by mating a haploid strain with a diploid strain of the opposite mating type (haploid *Mata* with diploid *Matα*/*Matα* or haploid *Matα* with diploid *Mata*/*Mata*).

### Genome modifications

#### Mating-type switch

The diploid strains *Mata*/*Mata* and *Matα*/*Matα* were generated by switching one of the mating types of parent diploid strains *Mata*/*Matα.* The mating-type switch was done via two-step gene replacement strategies. Two different plasmids carrying the *URA3* marker and either the *Matα* allele (pSC9 = OPL233) or the *Mata* allele (pSC11 = OPL234) were used [34]. The diploid strains *Mata*/*Matα* were originally transformed by inserting the plasmids digested by EcoRI at the mating type locus. After transformation, *URA3*+ clones were selected. After growing the clones overnight on non-selective media (YPD), the revertant *ura-* clones were selected by using 5’FOA plate that is toxic for *URA*+ cells. Finally, the mating type of diploid *ura-* clones were tested.

#### Heterozygous *CEN1-GFP* dots

An array of 256 lacO operator sites on plasmid pJN2 was integrated near the *CEN1* locus (coordinates 153583–154854). *lacI-GFP* fusions under the control of *P_CYC1_* was also expressed in this strain to visualize the location of the *lacO* operator.

#### Gene modifications

PCR-based methods were used to create complete deletions of ORFs (*mps1::kanMX, mad2::NAT)* [35, 36]*. mps1-R170S::his5* strains were generated previously [20].

##### SPC29-GFP

*SPC29* was tagged with GFP at its C-terminal at its original locus using one-step PCR method and the plasmid OPL436 (pFA6a-link-yoEGFP-Kan, http://www.addgene.org/44900).

##### mps1-aid*

*MPS1* gene was fused with an auxin-inducible degron tag at its C-terminal at its original gene locus, using the one-step PCR method, as described previously [21]. Using two plasmids (pKan–AID*–9myc and pNat–AID*–9myc), we generated two different versions of Mps1 C-terminal tagged: *MPS1-AID*-9myc::NAT* and *MPS1-AID*-9myc::KAN*.

##### *HphNT1-RS1-CEN3-RS1* and *CEN3-RS2-HIS3MX6*

*HPH*+ and *HIS*+ markers were inserted at close proximity to the centromere of chromosome 3 to exclude among the *ura-* revertant the one coming from chromosome loss.

##### F-Box protein AFB2

As an auxin receptor, we used *AFB2*, which promote an enhanced degradation in a yeast-based system compare to classically used *TIR1* [37]. We placed *AFB2* expression under the control of either the *GPD1* (*P_GPD_-AFB2-LEU2*) or *CUP1* promoter (*P_CUP1_-AFB2-HIS*).

###### Plasmid shuffle

A plasmid containing a copy of *MPS1-as1* and the *TRP1* yeast marker (OPL77 *=* pRS314-*MPS1-as1*, a gift from Mark Winey) was added to the cells to keep them alive when having a null allele of *MPS1* or prevent the accumulation of suppressor mutations when having the *mps1-R170S* allele. The shuffling plasmid was counter-selected by placing the cells on media containing the compound 5-Fluoroanthranilic acid (FAA). This compound is an antimetabolite for the tryptophan pathway in yeast, and is toxic by virtue of its antimetabolic conversion to 5-fluorotryptophan [38].

### Fluorescence microscopy

#### Long-term time-lapse microscopy

Images were acquired in 2-2.5min intervals for 6–8 h with exposure times of 100-200ms according to fluorescence intensity. Cells were imaged using ONIX Microfluidic Perfusion System from CellAsic with a flow rate of 5 psi (http://www.emdmillipore.com/US/en/life-science-research/cell-culture-systems/cellASIC-live-cell-analysis/microfluidic-plates/). Cells were incubated at 30°C to mid–log phase and loaded into the chamber for haploid, diploid and triploid cells. Tetraploid cells were directly loaded into the chamber. Haploid cells were loaded on Y04C plates whereas diploid, triploid and tetraploid cells were loaded on Y04D or Y04E plates. For haploid, diploid and triploid cells, time-lapse imaging was started 30-60mn after the loading into the chamber and was performed at 30 °C. For tetraploid cells, time-lapse imaging was started 3h30-4h after loading into the chamber and was performed at 30 °C.

For the time-lapse imaging following the behavior of the spindle, a marker for SPBs (*SPC29-GFP)* was used. Images were originally collected with a Nikon Eclipse TE2000-E equipped with the Perfect Focus system, a Roper CoolSNAP HQ2 camera automated stage, an X-cite series 120 illuminator (EXFO) and NIS software. Images were later collected with a Nikon Eclipse Ti2, a Hamamatsu ORCA-Flash4.0 V3 Digital CMOS C13440-20CU camera, a Lumencor SPECTRA X light engine and NIS software. Time-lapse microscopy was analyzed for cell cycle duration. The formation of the spindle was defined as the time when two separated SPBs can be easily observed. Anaphase onset was defined as the time when a rapid elongation of the spindle was observed. Data were graphed in Prism (GraphPad Software) and the significance was calculated using the Unpaired t test.

#### High speed time-lapse microscopy

Time-lapse imaging (every 5-10 second for 5-10 minutes) were collected using a Roper CoolSNAP HQ2 camera on a Zeiss Axio Imager 7.1 microscope fitted with a 100×, NA1.4 plan-Apo objective (Carl Zeiss MicroImaging), an X-cite series 120 illuminator (EXFO). Cells were incubated at 30°C to mid–log phase. Dividing cells were concentrated, spread across polyethyleneimine-treated coverslips, then covered with a thin 1% agarose pad to anchor the cells to the coverslip [39]. The coverslip was then inverted over a silicone rubber gasket attached to a glass slide. The measurements of the spindle length were all done at the beginning of the movies (t=0mn). For the analysis of type of *CEN1-GFP* attachment on bipolar spindle, the position of the centromeres was estimated at the beginning and at the end (t=5mn) of the movie. The centromeres were defined as monopolar when sister chromatids remain un-separated at close proximity (<0.5μm) to the SPBs for at least three consecutive frames. Centromeres were considered to be bipolar with un-separated sister chromatid when they remain at a central position on the spindle and the sister chromatids appears as a single dot for at least three consecutive frames. Centromeres were considered to be bipolar separated when the sister chromatids were observed at two separated dots for at least three consecutive frames. Cells were defined as converted when able to switch from a monopolar attachment at t=0mn to a bipolar attachment at t=5mn. Cells were defined as unconverted when observed as monopolar attachment at t=0mn and t=5mn. For the analysis of type of *CEN1-GFP* attachment on pre-anaphase spindle, the anaphase onset (t=0s) was initially determined by observing a rapid elongation of the spindle. The pre-anaphase attachment was determined by observing the type of attachment one minute before this event (t=−60s). The centromeres were defined as monopolar when the sister chromatids remain un-separated and at close proximity (<0.5μm) to the SPBs for at least three consecutive frames. Centromeres were considered to be bipolar with low stretching when the distance between the two sister chromatids was under 0.5μm. Centromeres were considered to be bipolar with high stretching when the distance between the two sister chromatids was over 0.5μm.

## Abbreviations

SPB: spindle pole body

## Acknowledgements

We thank Dean Dawson for critically reading the manuscript and providing strains and reagents. We thank current and past laboratory members for reagents, and discussions of our work. AS, DH, MG and JH contributed to this project as participants in the OMRF Fleming Scholars Program for undergraduate summer research.

---

**S1 Fig.**
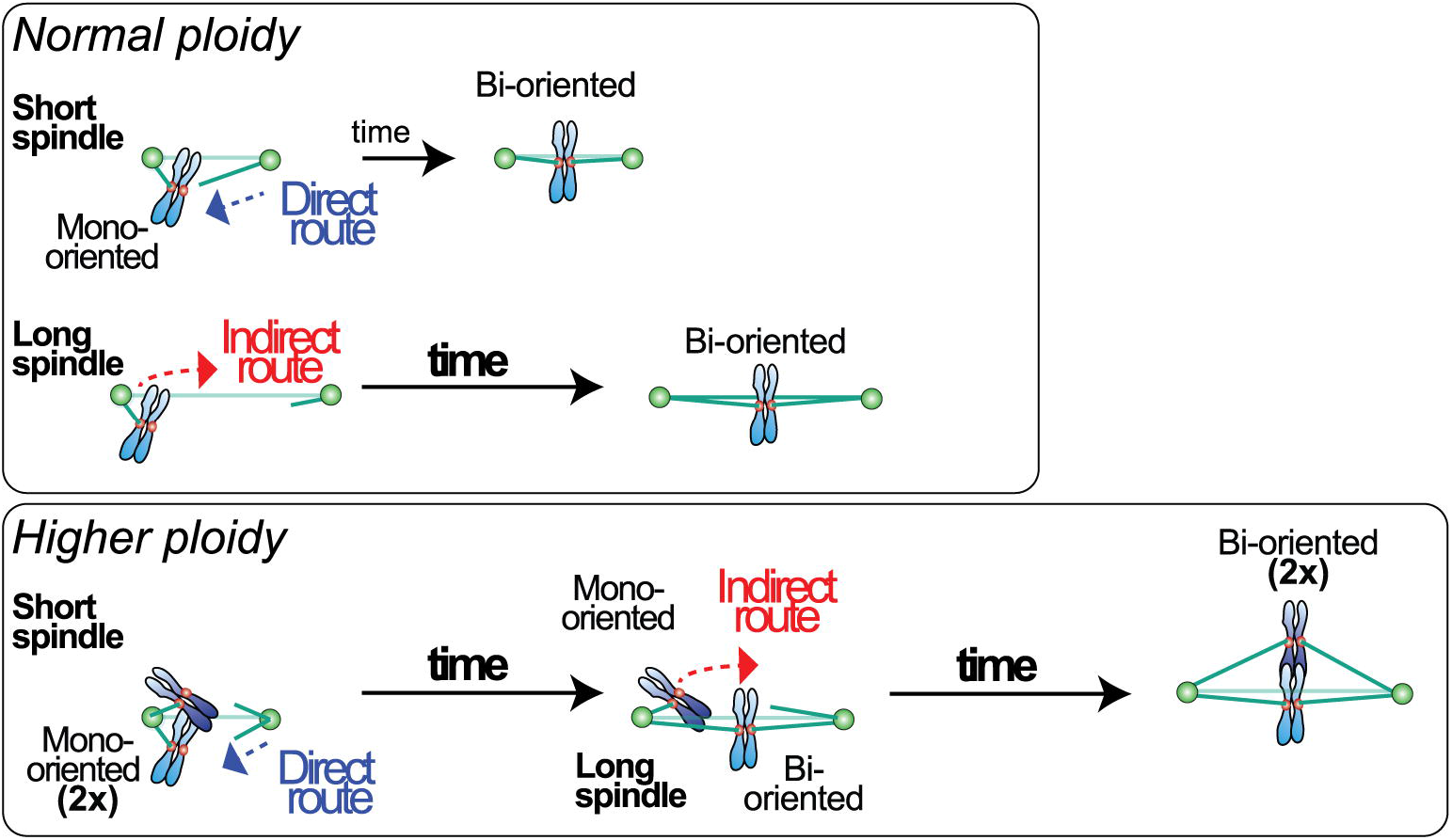
Effect of *mps1* mutations on cell viability depending of the ploidy. Ten-fold serial-dilution growth assays. All the indicated strains for the different ploidies (Haploid=1n, Diploid=2n, Triploid=3n, Tetraploid=4n) are carrying a shuffling plasmid expressing *MPS1-as1* and the yeast marker *TRP1*. Cells were originally grown in CM-TRP media and later spotted on the indicated plates. The CM-TRP plates were used to select yeast carrying the shuffling plasmid with the extra copy of Mps1 whereas the FAA plates were used to counter-select it. Tetraploid cells with four copies of either *MPS1* allele (*MPS1-4x = MPS1/MPS1/MPS1/MPS1*) or *mps1-R170S* allele (*mps1-R170S-4x = mps1-R170S/mps1-R170S/mps1-R170S/ mps1-R170S)* or *mps1Δ (MPS1-0x = mps1Δ/mps1Δ/mps1Δ/mps1Δ)* were used in this assay.

**S2 Fig. Viability assay on *mps1-AID*\* tetraploid cells.** Ten-fold serial-dilution growth assays for tetraploid cells. All the tetraploid cells of the indicated genotype are expressing the F-protein *AFB2*. The cells were originally grown in YPAD media and later spotted on the indicated plates. The YPAD plates with Auxin (0.5mM) were used to induce the degradation of the Mps1 tagged with the AID* degron system.

**S3 Fig. Cell division and SPB duplication assays.** (A-C). All cells (haploid and tetraploid) are carrying a single copy of Spc29-GFP. (A) Representative pictures of wild-type and *mad2* mutant tetraploid cells growing inside the microfluidic chamber are shown. Indicated time is in minutes. Bars 10 μm. (B) The cell viability shown in Fig. 2A was estimated by following the behavior of the progeny during the 1^st^ two cell cycles. We evaluated the proportion of non-growing/dead cells after each cell division. In the representative *MPS1-1x* mutant tetraploid cells shown, we observed two viable cells after the first cell division (2 of 2). In the second round of cell division (cell division #2 and #3), we noticed that half of the progeny (2 of 4) were not viable as shown on the representative pictures. Indicated time is in minutes. Bars 10 μm. (C) The ability of cells to duplicate their SPBs was estimated after bud formation. The proportion of cells able to duplicate is shown. * = p < 0.05, (Fischer’s exact t test).

**S4 Fig. Conversion from monopolar to bipolar attachment in wild-type tetraploid cells.** The ability of individual tetraploid wild-type cells to convert monopolar to bipolar attachment in 5 minutes was evaluated depending of the spindle size. The proportion of cell able to convert in this interval depending of the indicated spindle length is shown. At least 19 cells were counted per categories. ns=non-significant (Fischer’s exact t test).

**S1 Table. Strains used in this study.**

**S2 Table. Diploid parent strain list.**

**S3 Table. Haploid parent strain list.**

