## Supplementary figures and images for "Mps1 expression is critical for chromosome orientation and segregation in yeast with high ploidy"

### Supplemental Figure 1

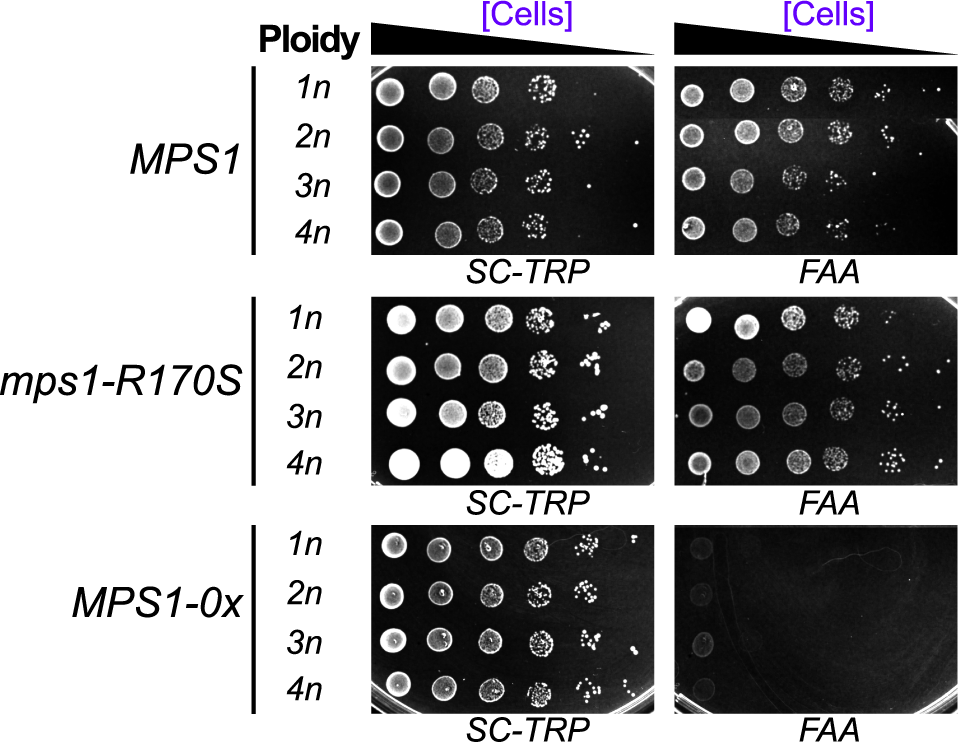

### Supplemental Figure 2

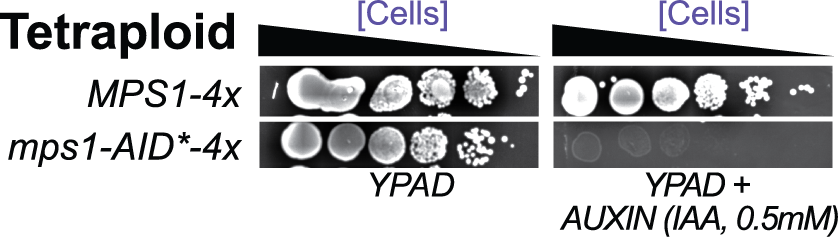

### Supplemental Figure 3

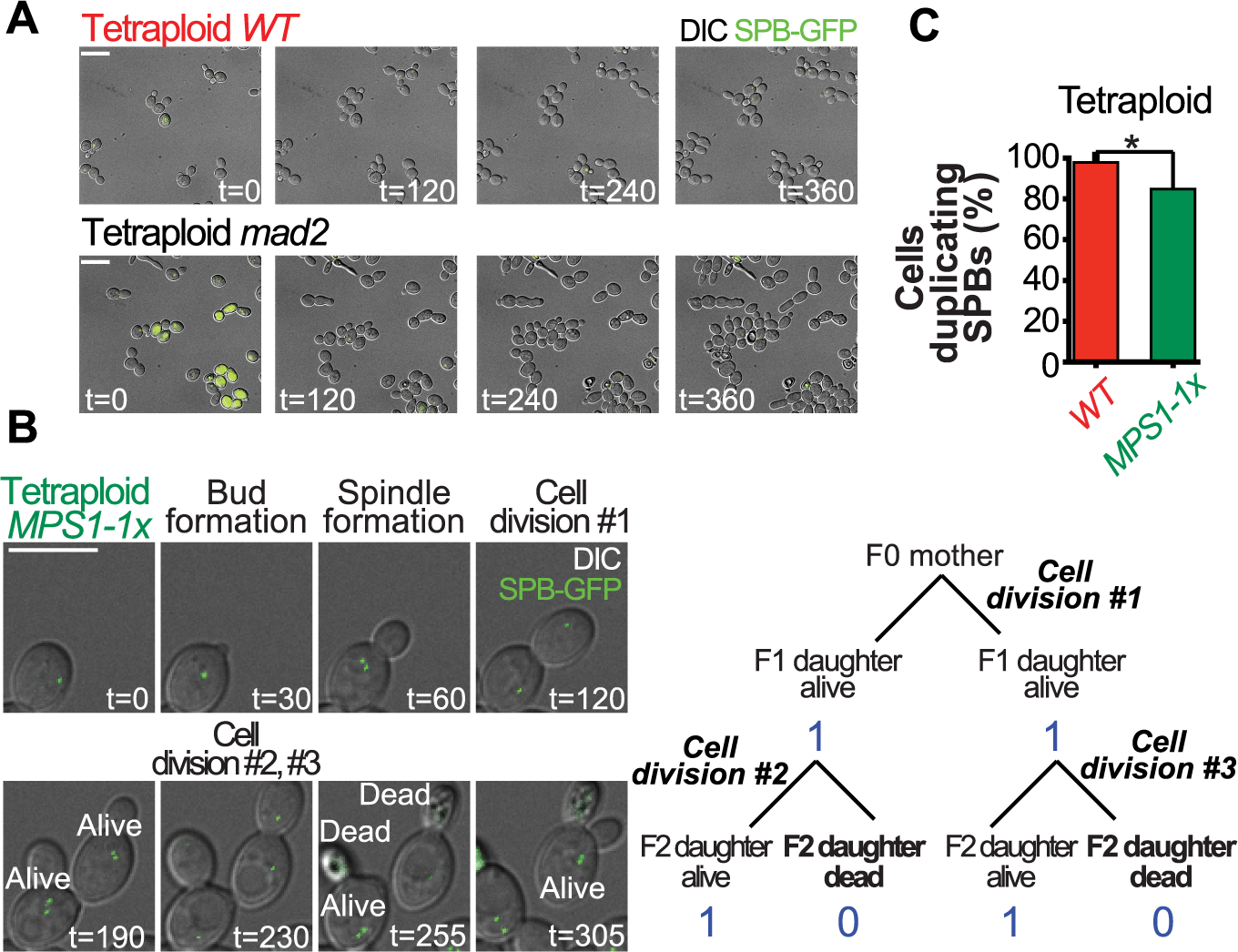

### Supplemental Figure 4

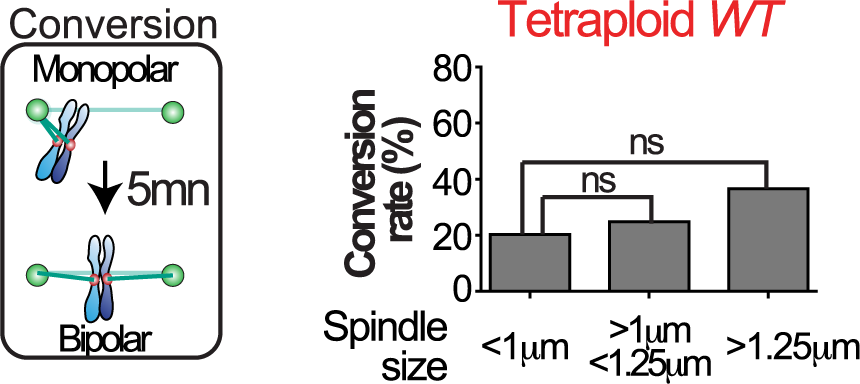
