## Supplemental Table 1 for "Mps1 expression is critical for chromosome orientation and segregation in yeast with high ploidy"

| **Figure** | **Ploidy and**  **Genotype** | **Strains** | **Parents strain**  (Haploid or diploid) |
| --- | --- | --- | --- |
| **Fig. 1B** | Tetraploid *MPS1-4x* | O503 | O485 * O486 |
|  | Tetraploid *MPS1-2x* | O549 | O485 * O521 |
| **Fig. 1C** | Tetraploid *mps1-R170S-4x* | O519 | O494 * O496 |
|  | Tetraploid *mps1-R170S-2x* | O560 | O494 * O524 |
|  | Tetraploid *MPS1-0x* | O527 | O521 * O524 |
| **Fig. 1A** | Diploid *MPS1-2x* | O485 |  |
|  | Diploid *MPS1-1x* | O489 |  |
|  | Diploid *mps1-R170S-1x* | O497 |  |
|  | Diploid *MPS1-0x* | O501 |  |
| **Fig. 2A** | Haploid *WT* | X3177 |  |
| **Fig. 2A, 3A-B, S3** | Tetraploid *WT* | 4nDH111  4nRM124 | O753 * O630,  O750 * O627 |
|  | Tetraploid *mad2* | 4nDH113  4nRM126 | O849 * O576,  O852 * O573 |
|  | Tetraploid *MPS1-1x* | 4nDH115  4nDH116 | O853 * O860,  O856 * O857 |
| **Fig. 2B-C, 3** | Haploid *WT* | X3177, X3178 |  |
| **Fig. 2C** | Haploid *mad2* | X3469, X3470 |  |
|  | Haploid *MPS1-1x* | X3473, X3474 |  |
|  | Diploid *WT* | O753, O750, O749, O754 |  |
|  | Diploid *mad2* | O849, O852, O850 |  |
|  | Diploid *MPS1-1x* | DDH3713, DRM4019 DAS4047, DRM4020, DAS4047 | X3473 * X2834  X3474 * X2833 |
| **Fig. 2B-C** | Triploid *WT* | 3nDH21  3nRM33  3nRM34 | O753 * X206  O754 * X206  O749 * X55 |
| **Fig. 2C** | Triploid *mad2* | 3nDH23, 3nRM35  3nDH24 | O849 * X259  O852 * X258 |
| **Fig. 2C** | Triploid *MPS1-1x* | 3nDH25, 3nRM37  3nRM38 | O853 * X2834  O856 * X2833 |
| **Fig. 2B-C, 3D** | Tetraploid *WT* | 4nRM123  4nAS135  4nAS136 | O753 * O630  O754 * O629  O750 * O627 |
| **Fig. 2C** | Tetraploid *mad2* | 4nRM125  4nAS137  4nAS138 | O849 * O576  O850 * O575  O852 * O573 |
| **Fig. 2B-C** | Tetraploid *MPS1-1x* | 4nRM127  4nRM128  4nAS140 | O853 * O860  O856 * O857  O855 * O858 |

**S1 Table: Strains used in this study.**

| **Figure** | **Ploidy and**  **Genotype** | **Strains** | **Parents strain**  (Haploid or diploid) |
| --- | --- | --- | --- |
| **Fig. 4-5, S4** | Haploid *WT* | X3493, X3494 |  |
|  | Diploid *WT* | DAS3717, DAS4707 | X3177 * X754 |
|  |  | DAS3718 | X3178 * X753 |
|  | Tetraploid *WT* | 4nAS117 | O753 * O904 |
|  |  | 4nAS118, 4nAS142 | O750 * O903 |
|  |  | 4nAS131 | O754 * O904 |
|  |  | 4nAS132 | O749 * O903 |
|  | Tetraploid *MPS1-1x* | 4nAS119, 4nAS143 | O853 * O907 |
|  |  | 4nAS120 | O856 * O905 |
|  |  | 4nAS133 | O854 * O908 |
|  |  | 4nAS134 | O855 * O906 |
| **Fig. S1** | Haploid *WT* | Y2179 |  |
|  | Diploid *WT* | O487 |  |
|  | Triploid *WT* | O533 | O487 * Y170 |
|  | Tetraploid *WT* | O539 | O487 * O488 |
|  | Haploid *mps1-R170S* | Y2183 |  |
|  | Diploid *mps1-R170S* | O495 |  |
|  | Triploid *mps1-R170S* | O526 | O484 * Y2183 |
|  | Tetraploid *mps1-R170S* | O518 | O495 * O496 |
|  | Haploid *MPS1-0X* | Y2181 |  |
|  | Diploid *MPS1-0X* | O529 |  |
|  | Triploid *MPS1-0X* | O536 | Y2181 * O524 |
|  | Tetraploid *MPS1-0X* | O541 | O523 * O524 |
| **Fig. S2** | Tetraploid *MPS1-4X* | O655 | O631 * O633 |
|  | Tetraploid *mps1AID*-4X* | O755 | O635 * O638 |
