## Supplemental Table 2 for "Mps1 expression is critical for chromosome orientation and segregation in yeast with high ploidy"

**S2 Table: Diploid parent strain list.**

***XX strain***: *ura3-13, trp1-Δ63, his3-Δ1, leu2-?, met13-d, tyr1-1, lys2-1, can1-R*

***YY strain***: *ura3-1, trp1-Δ63, his3-Δ1, leu2-?, met13-c, tyr1-2, lys2-2, cyh2-1*

| O485 | ***XX strain,*** *MATa/MATa, CEN-plasmid pRS314-mps1-as1* |
| --- | --- |
| O486 | ***XX strain,*** *MAT∝/MAT∝, CEN-plasmid pRS314-mps1-as1* |
| O521 | ***XX strain,*** *MAT∝/MAT∝, mps1∆::KANMX/mps1∆::KANMX, CEN-plasmid pRS314-mps1-as1* |
| O494 | ***XX strain,*** *MAT∝/MAT∝, mps1-R170S::his5/mps1-R170S::his5, CEN-plasmid pRS314-mps1-as1* |
| O496 | ***XX strain,*** *MATa/MATa, mps1-R170S::his5/mps1-R170S::his5, CEN-plasmid pRS314-mps1-as1* |
| O524 | ***XX strain,*** *MATa/MATa, mps1∆::KANMX/mps1∆::KANMX, CEN-plasmid pRS314-mps1-as1* |
| O489 | ***XX strain,*** *MATa/MATa, mps1∆::KANMX/MPS1, CEN-plasmid pRS314-mps1-as1* |
| O497 | ***XX strain,*** *MAT∝/MAT∝, mps1-R170S::his5/mps1∆::KANMX, CEN-plasmid pRS314-mps1-as1-TRP1* |
| O501 | ***XX strain,*** *MAT∝/MAT∝, mps1∆::KANMX/mps1∆::KANMX CEN-plasmid pRS314-mps1-as1-TRP1,* |
| O753, O754 | ***XX strain,*** *MATa/MATa, CEN3-RS2-HIS3MX6/HphNT1-RS1-CEN3, SPC29-yoEGFP Kan/+* |
| O749, O750 | ***XX strain,*** *MAT∝/MAT∝, CEN3-RS2-HIS3MX6/HphNT1-RS1-CEN3, SPC29-yoEGFP Kan/+* |
| O627 | ***XX strain,*** *MATa/MATa CEN3-RS2-HIS3MX6/HphNT1-RS1-CEN3* |
| O629, O630 | ***XX strain,*** *MAT∝/MAT∝, CEN3-RS2-HIS3MX6/HphNT1-RS1-CEN3* |
| O849, O850 | ***XX strain,*** *MATa/MATa, CEN3-RS2-HIS3MX6/HphNT1-RS1-CEN3, SPC29-yoEGFP Kan/+, mad2::NATMX6/mad2::NATMX6* |
| O852 | ***XX strain,*** *MAT∝/MAT∝, CEN3-RS2-HIS3MX6/HphNT1-RS1-CEN3, SPC29-yoEGFP Kan/+, mad2::NATMX6/mad2::NATMX6* |
| O573 | ***XX strain,*** *MATa/MATa, mad2::NATMX6/mad2::NATMX6* |
| O575, O576 | ***XX strain,*** *MAT∝/MAT∝, mad2::NATMX6/mad2::NATMX6* |
| O853, O854 | ***XX strain,*** *MATa/MATa, tyr1::[HIS5 pCUP1-AFB2]/tyr1-1, CEN3-RS1-hphNT1/+, SPC29-yoEGFP Kan/+, mps1Δ::KANMX4/+* |
| O855, O856 | ***XX strain,*** *MAT∝/MAT∝, tyr1::[HIS5 pCUP1-AFB2]/tyr1-1, CEN3-RS1-hphNT1/+, SPC29-yoEGFP Kan/+, mps1Δ::KANMX4/+* |
| O860 | ***XX strain,*** *MAT∝/MAT∝, CEN3-RS2-HIS3MX6/HphNT1-RS1-CEN3, MPS1-AID*-9myc-natNT2/mps1Δ::KANMX4* |
| O857, O858 | ***XX strain,*** *MATa/MATa, CEN3-RS2-HIS3MX6/HphNT1-RS1-CEN3, MPS1-AID*-9myc-natNT2/mps1Δ::KANMX4* |
| O903 | ***XX strain,*** *MATa/MATa, CEN3-RS2-HIS3MX6/HphNT1-RS1-CEN3, lys2::pLL1[PCYC1-GFP-lacI LYS2]/lys2-2, CEN1::pJN2[lacO256 LEU2]/+* |
| O904 | ***XX strain,*** *MAT∝/MAT∝, CEN3-RS2-HIS3MX6/HphNT1-RS1-CEN3, lys2::pLL1[PCYC1-GFP-lacI LYS2]/lys2-2, CEN1::pJN2[lacO256 LEU2]/+* |

| O905, O906 | ***XX strain,*** *MATa/MATa, CEN3-RS2-HIS3MX6/HphNT1-RS1-CEN3, lys2::pLL1[PCYC1-GFP-lacI LYS2]/lys2-2, CEN1::pJN2[lacO256 LEU2]/+, MPS1-AID*-9myc-natNT2/mps1Δ::KANMX4, CEN1::pJN2[lacO256 LEU2]/+* |
| --- | --- |
| O907, O908 | ***XX strain,*** *MAT∝/MAT∝, CEN3-RS2-HIS3MX6/HphNT1-RS1-CEN3, lys2::pLL1[PCYC1-GFP-lacI LYS2]/lys2-2, CEN1::pJN2[lacO256 LEU2]/+, MPS1-AID*-9myc-natNT2/mps1Δ::KANMX4, CEN1::pJN2[lacO256 LEU2]/+* |
| O487 | ***YY strain,*** *MAT∝/MAT∝, CEN-plasmid pRS314-mps1-as1* |
| O488 | ***YY strain,*** *MATa/MATa, CEN-plasmid pRS314-mps1-as1* |
| O495 | ***YY strain,*** *MAT∝/MAT∝, mps1-R170S::his5/mps1-R170S::his5,* *CEN-plasmid pRS314-mps1-as1* |
| O484 | ***YY strain,*** *MATa/MATa, mps1-R170S::his5/mps1-R170S::his5* |
| O496 | ***YY strain,*** *MATa/MATa, mps1-R170S::his5/mps1-R170S::his5,* *CEN-plasmid pRS314-mps1-as1* |
| O529 | ***YY strain,*** *MATa/MAT∝,* *mps1∆::KANMX/mps1∆::KANMX, CEN-plasmid pRS314-mps1-as-1-TRP1* |
| O523 | ***YY strain,*** *MAT∝/MAT∝, mps1∆::KANMX/mps1∆::KANMX, CEN-plasmid pRS314-mps1-as-1-TRP1* |
| O524 | ***YY strain,*** *MATa/MATa,* *mps1∆::KANMX/mps1∆::KANMX, CEN-plasmid pRS314-mps1-as-1-TRP1* |
| O631 | ***XX strain,*** *MATa/MATa, CEN3-RS2-HIS3MX6/HphNT1-RS1-CEN3, leu2::[OPL281:LEU2, PGPD-AFB2]/leu2::[OPL281:LEU2, PGPD-AFB2]* |
| O633 | ***XX strain,*** *MAT∝/MAT∝,, CEN3-RS2-HIS3MX6/HphNT1-RS1-CEN3, leu2::[OPL281:LEU2, PGPD-AFB2]/leu2::[OPL281:LEU2, PGPD-AFB2]* |
| O635 | ***XX strain,*** *MATa/MATa, CEN3-RS2-HIS3MX6/HphNT1-RS1-CEN3, leu2::[OPL281:LEU2, PGPD-AFB2]/leu2::[OPL281:LEU2, PGPD-AFB2], MPS1-AID*-9Myc-KAN/MPS1-AID*-9Myc-KAN* |
| O638 | ***XX strain,*** *MAT∝/MAT∝,, CEN3-RS2-HIS3MX6/HphNT1-RS1-CEN3, leu2::[OPL281:LEU2, PGPD-AFB2]/leu2::[OPL281:LEU2, PGPD-AFB2], MPS1-AID*-9Myc-KAN/MPS1-AID*-9Myc-KAN* |
