## Supplemental Table 3 for "Mps1 expression is critical for chromosome orientation and segregation in yeast with high ploidy"

**S3 Table: Haploid parent strain list.**

| X3177 | *MATa, ura3-13, trp1-Δ63, leu2-?, tyr1-1, lys2-1, met13-d, can1-R, SPC29-yoEGFP Kan, CEN3-RS2-HIS3MX6* |
| --- | --- |
| X3178 | *MAT∝, ura3-13, trp1-Δ63, leu2-?, tyr1-1, lys2-1, met13-d, can1-R, SPC29-yoEGFP Kan, CEN3-RS2-HIS3MX6* |
| X3469 | *MATa, ura3-13, trp1-Δ63, leu2-?, tyr1-1, lys2-1, met13-d, can1-R, SPC29-yoEGFP Kan, CEN3-RS2-HIS3MX6, mad2::NATMX6* |
| X3470 | *MAT∝, ura3-13, trp1-Δ63, leu2-?, tyr1-1, lys2-1, met13-d, can1-R, SPC29-yoEGFP Kan, CEN3-RS2-HIS3MX6, mad2::NATMX6* |
| X3473 | *MATa, ura3-13, trp1-Δ63, leu2-?, tyr1::[HIS5 pCUP1-AFB2], lys2-1, met13-d, can1-R, SPC29-yoEGFP Kan* |
| X3474 | *MAT∝, ura3-13, trp1-Δ63, leu2-?, tyr1::[HIS5 pCUP1-AFB2], lys2-1, met13-d, can1-R, SPC29-yoEGFP Kan* |
| X2833 | *MATa, ura3-13, trp1-Δ63, his3-Δ1, leu2-?, met13-d, tyr1-1, lys2-1, can1-R, HphHNT1-RS1-CEN3, MPS1-AID*-9Myc-NAT* |
| X2834 | *MAT∝, ura3-13, trp1-Δ63, his3-Δ1, leu2-?, met13-d, tyr1-1, lys2-1, can1-R, HphHNT1-RS1-CEN3, MPS1-AID*-9Myc-NAT* |
| X55 | *MATa, ura3-13, trp1-Δ63, his3-Δ1, leu2-?, met13-d, tyr1-1, lys2-1, can1-R* |
| X206 | *MAT∝, ura3-13, trp1-Δ63, leu2, tyr1-1, lys2-1, met13-d, can1-R, his3-Δ1* |
| X258 | *MATa, ura3-13, trp1-Δ63, leu2, tyr1-1, lys2-1, met13-d, can1-R, his3-Δ1, mad2::NATMX6* |
| X259 | *MAT∝, ura3-13, trp1-Δ63, leu2, tyr1-1, lys2-1, met13-d, can1-R, his3-Δ1, mad2::NATMX6* |
| X3493 | *MATa, ura3-13, trp1-Δ63, his3-Δ1, leu2-?, met13-d, tyr1-1, lys2::pLL1[PCYC1-GFP-lacI LYS2], can1-R, CEN1::pJN2[lacO256 LEU2], SPC29-yoEGFP Kan* |
| X3494 | *MATa, ura3-13, trp1-Δ63, his3-Δ1, leu2-?, met13-d, tyr1-1, lys2::pLL1[PCYC1-GFP-lacI LYS2], can1-R, CEN1::pJN2[lacO256 LEU2], SPC29-yoEGFP Kan* |
| X753 | *MATa, can1-R, leu2, lys2-1::pLL1[PCYC1-GFP-lacI LYS2], met13-d, trp1-Δ63, tyr1-1, ura3-13, his3-Δ1, CEN1::pJN2[lacO256 LEU2]* |
| X754 | *MAT∝, can1-R, leu2, lys2-1::pLL1[PCYC1-GFP-lacI LYS2], met13-d, trp1-Δ63, tyr1-1, ura3-13, his3-Δ1, CEN1::pJN2[lacO256 LEU2]* |
| Y2179 | *MAT∝, leu2-?, lys2-2, met13-c, tyr1-2, ura3-1, trp1-Δ63, cyh2-1, his3-Δ1, CEN-plasmid pRS314-mps1-as1-TRP1* |
| Y170 | *MATa, leu2-?, lys2-2, met13-c, tyr1-2, ura3-1, trp1-Δ63, cyh2-1, his3-Δ1* |
| Y2183 | *MAT∝, leu2-?, lys2-2, met13-c, tyr1-2, ura3-1, trp1-Δ63, cyh2-1, his3-Δ1, CEN-plasmid pRS314-mps1-as1-TRP1, mps1-R170S::his5* |
| Y2181 | *MAT∝, leu2-?, lys2-2, met13-c, tyr1-2, ura3-1, trp1-Δ63, cyh2-1, his3-Δ1, CEN-plasmid pRS314-mps1-as1-TRP1, mps1∆::KANMX* |
